# 40 minutes RT-qPCR Assay for Screening Spike N501Y and HV69-70del Mutations

**DOI:** 10.1101/2021.01.26.428302

**Authors:** Gulay Korukluoglu, Mustafa Kolukirik, Fatma Bayrakdar, Gozde Girgin Ozgumus, Ayse Basak Altas, Yasemin Cosgun, Canan Zohre Ketre Kolukirik

**Affiliations:** WHO National Influenza Center and Virology Reference Laboratory of Ministry of Health Turkey, Ankara, Turkey; Bioeksen R&D Technologies Limited, Istanbul Technical University Teknokent, Istanbul, Turkey

**Keywords:** SARS-CoV-2, VOC-202012/01, B.1.1.7, 501Y.V2, B.1.351, P.1, Spike N501Y, Spike HV69-70del, Mutation, Variant

## Abstract

A one-step reverse transcription and real-time PCR (RT-qPCR) test was developed for rapid screening (40 minutes) of the Spike N501Y and HV69-70del mutations in SARS-CoV-2 positive samples. The test also targets a conserved region of SARS-CoV-2 Orf1ab as an internal control. The samples containing both the N501Y and HV69-70del mutations are concluded as VOC-202012/01 positive. Samples suspected to be positive for B.1.351 or P.1 are the N501Y positive and HV69-70del negative cases. Limit of detection (LOD) of the kit for Orf1ab target is 500 copies/mL, while that of the N501, Y501 and HV69-70del targets are 5000 copies/mL. The developed assay was applied to 165 clinical samples containing SARS-CoV-2 from 32 different lineages. The SARS-CoV-2 lineages were determined via the next-generation sequencing (NGS). The RT-qPCR results were in 100% agreement with the NGS results that 19 samples were N501Y and HV69-70del positive, 10 samples were N501Y positive and HV69-70del negative, 1 sample was N501Y negative and HV69-70del positive, and 135 samples were N501Y and HV69-70del negative. All the VOC-202012/01 positive samples were detected in people who have traveled from England to Turkey. The RT-qPCR test and the Sanger sequencing was further applied to 1000 SARS-CoV-2 positive clinical samples collected in Jan2021 from the 81 different provinces of Turkey. The RT-qPCR results were in 100% agreement with the Sanger sequencing results that 32 samples were N501Y positive and HV69-70del negative, 4 samples were N501Y negative and HV69-70del positive, 964 samples were N501Y and HV69-70del negative. The specificity of the 40 minutes RT-qPCR assay relative to the sequencing-based technologies is 100%. The developed assay is an advantageous tool for timely and representative estimation of the N501Y positive variants’ prevalence because it allows testing a much higher portion of the SARS-CoV-2 positives in much lower time compared to the sequencing-based technologies.

## INTRODUCTION

Increasing prevalence of the spike (S) N501Y mutation containing SARS-CoV-2 variants (Table 1) have significantly raised the concerns about the higher transmission rate (1) and the escape from neutralizing antibodies (2). VOC-202012/01 is correlated with a significant increase in the rate of COVID-19 infection in the United Kingdom (3). VOC-202012/01 could be up to 70% more transmissible than previous variants due to unusually large number of mutations in a single cluster, particularly in the S protein. A confirmed case of reinfection with SARS-CoV-2 with the second episode due to VOC-202012/01 was reported in the UK (4).

**Table 1.**
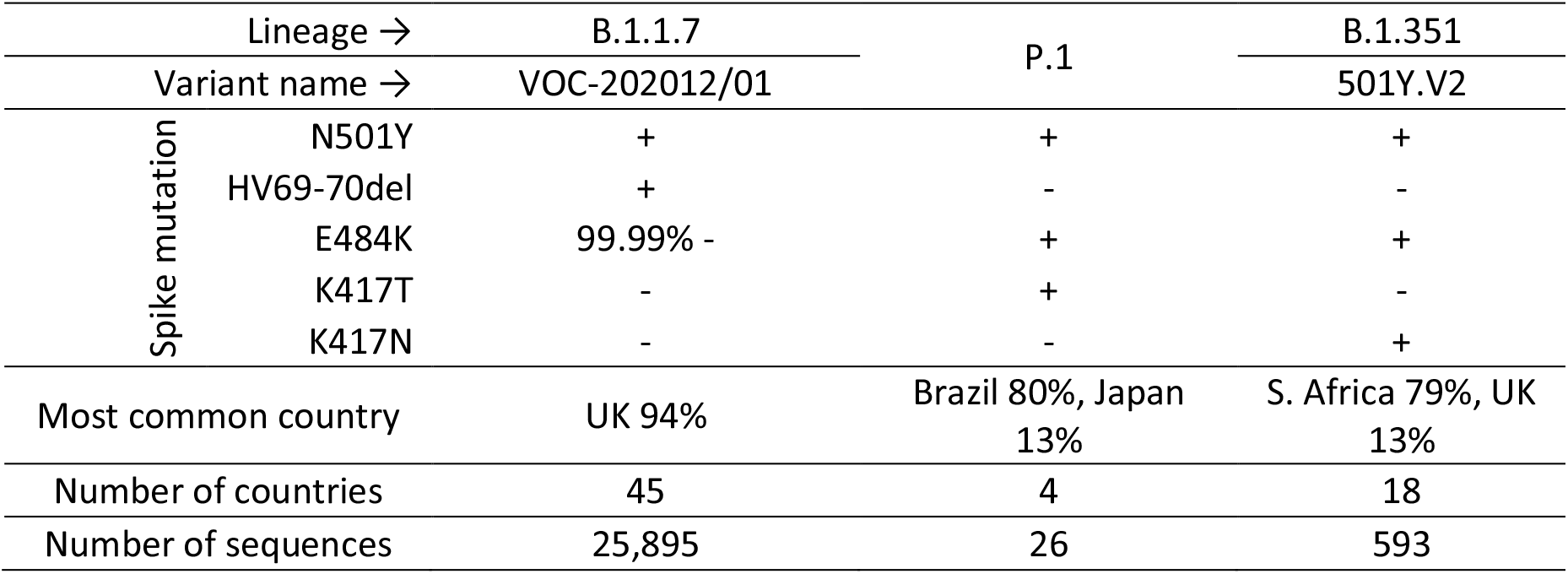
Statistics of the epidemiologically significant SARS-CoV-2 variants containing N501Y mutation (5).

| Lineage → |  | B.1.1.7 | P.1 | B.1.351 |
| --- | --- | --- | --- | --- |
| Variant name → |  | VOC-202012/01 |  | 501Y.V2 |
| Spike mutation | N501Y | + | + | + |
|  | HV69-70del | + | - | - |
|  | E484K | 99.99% - | + | + |
|  | K417T | - | + | - |
|  | K417N | - | - | + |
| Most common country |  | UK 94% | Brazil 80%, Japan 13% | S. Africa 79%, UK 13% |
| Number of countries |  | 45 | 4 | 18 |
| Number of sequences |  | 25,895 | 26 | 593 |

The N501Y is of particular importance among the other mutations of VOC-202012/01 (3). Mutations in C-terminal domain (CTD) (aa 333-527) of the S protein’s receptor-binding domain (RBD) is most likely to change receptor recognition properties of SARS-CoV-2 (6). N501Y sits in the receptor-binding motif and has been identified as increasing binding affinity to human ACE2 (7, 8).

Spike HV69-70del mutation of VOC-202012/01 is in the N terminal domain (NTD), has been identified in variants associated with immune escape in immunocompromised patients, and is responsible for a “dropout” in the S gene PCR target in a three-target (N, ORF1ab, S) commercial assay (3). 77% of the sequenced SARS-CoV-2 genomes containing HV69-70del is VOC-202012/01 (5). Therefore, screening only HV69-70del will not be decisive in regions where the VOC-202012/01 prevalence is low. The co-presence of N501Y and HV69-70del is a case only for VOC-202012/01 (5) and can be used for the monitoring regardless of the variant’s prevalence. In addition, the N501Y positive and HV69-70del negative cases are the possible candidates of infections with the variants 501Y.V2 or P.1 (Table 1).

In addition to the N501Y mutation, the 501Y.V2 and P.1 variants carry the S E484K and K417(T/N) mutations in the CTD of RBD. 9 of the 25,895 available VOC-202012/01 genomes contain the E484K mutation (5). Escape from neutralizing antibodies by the spike protein E484K variants was previously reported using a recombinant chimeric SARS-CoV-2 reporter virus (2). Molecular dynamic simulations have pointed out the possibility that the combination of E484K, K417N and N501Y mutations induces conformational change greater than N501Y mutant alone (9, 10).

Because all SARS-CoV-2 variants containing the N501Y are important for epidemiology, the timely and representative estimation of their prevalence by testing a much higher portion of the SARS-CoV-2 positives in much lower time compared to the sequencing-based technologies is necessary. In this study, a one-step reverse transcription and real-time PCR (RT-qPCR) test was developed for rapid screening of the Spike N501Y and HV69-70del mutations in SARS-CoV-2 positive samples.

## MATERIALS AND METHODS

### RT-qPCR Assay Targeting SARS-CoV-2 Spike N501Y and HV69-70del Mutations

The assay consumables were produced as a ready-to-use kit by Bioeksen R&D Technologies Limited, Turkey (Table 2). The N501Y detection with the kit is performed by comparing two different RT-qPCR reactions prepared using the “N501 Oligo Mix” and the “Y501 Oligo Mix”. A conserved region of ORF1ab is targeted in both reactions as an internal control. SARS-CoV-2 S protein mutation HV69-70del is also targeted in both reactions to differentiate VOC-202012/01 from the other N501Y containing variants (Table 1). The N501Y mutation detection is made by comparing the Cq value differences in the two reactions. The “Y501 Oligo Mix” must amplify Spike-501-target better (with lower Cq values) than the “N501 Oligo Mix” does to obtain the positive N501Y mutation result.

**Table 2.**
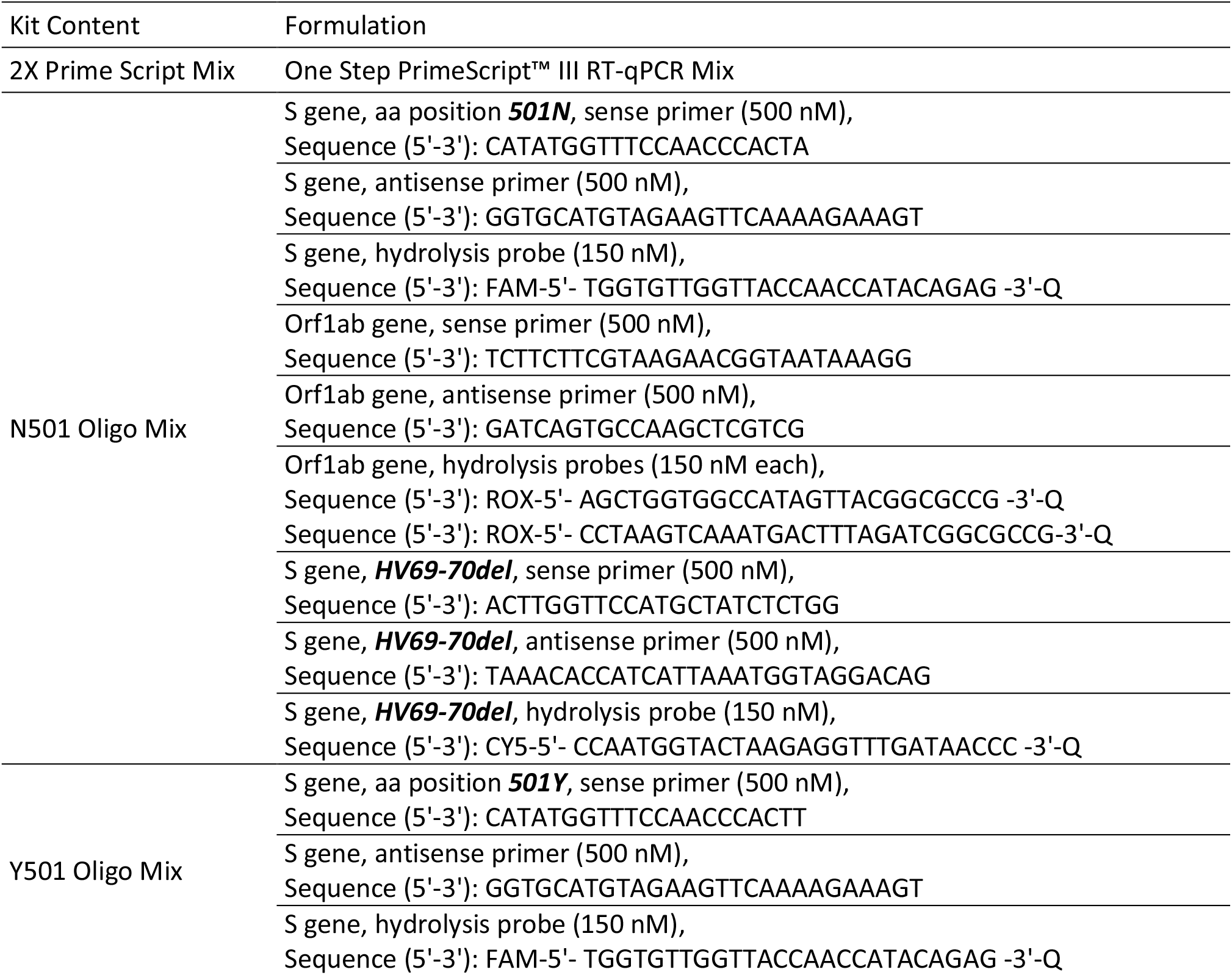
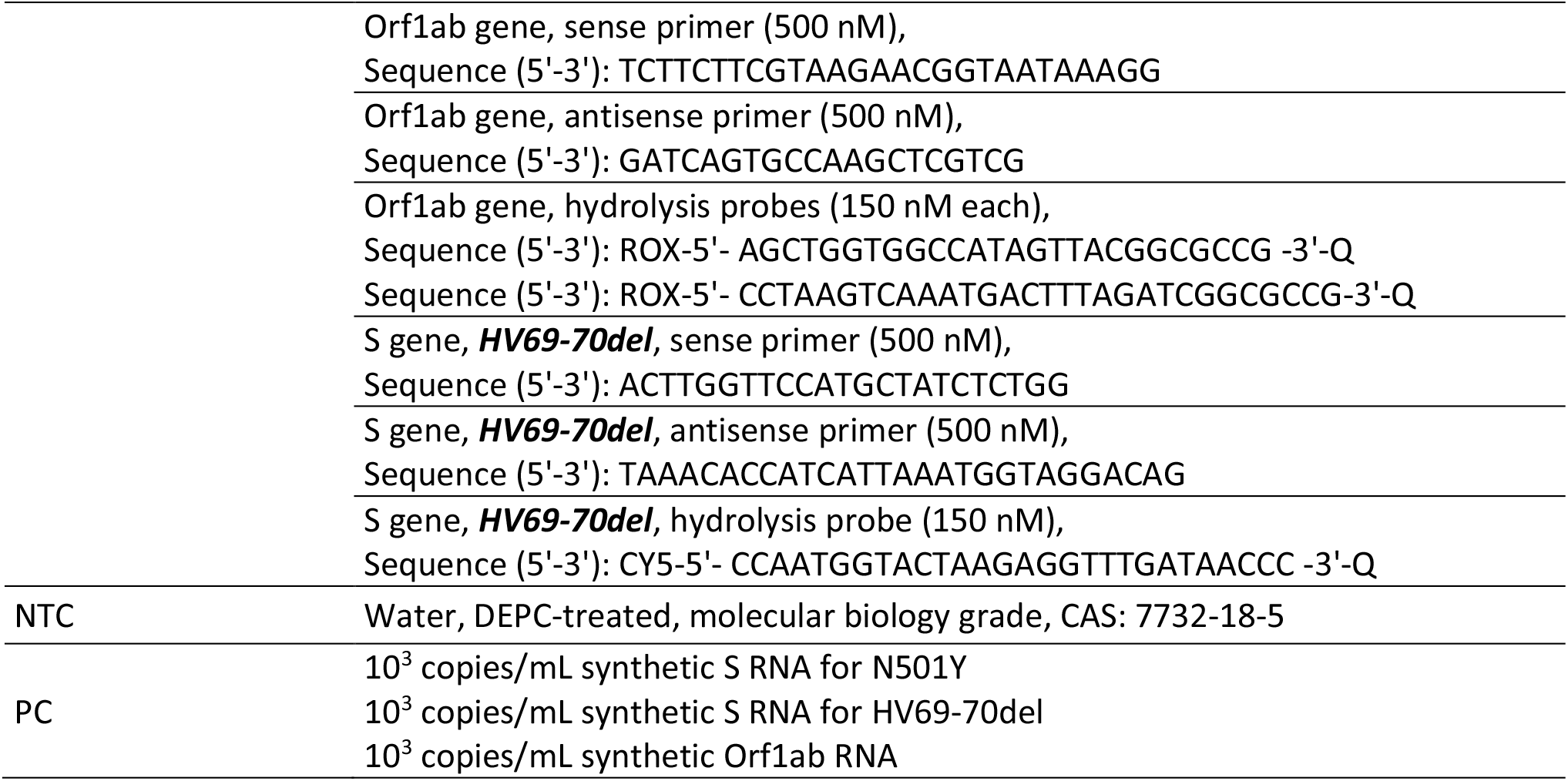
Contents of the “Bio-Speedy^®^ SARS-CoV-2 N501Y Mutation Detection Kit” (Cat No: BS-N501Y).

The SARS-CoV-2 genomes used for the oligonucleotide design (Table 3) reflect all the major lineages and the important variants emerged recently (11). The oligonucleotide sequences in Table 2 matches 100% with all their targets in Table 3. The oligonucleotide binding regions were also checked via CoV-GLUE in terms of the replacements, insertions and deletions (12).

**Table 3.**
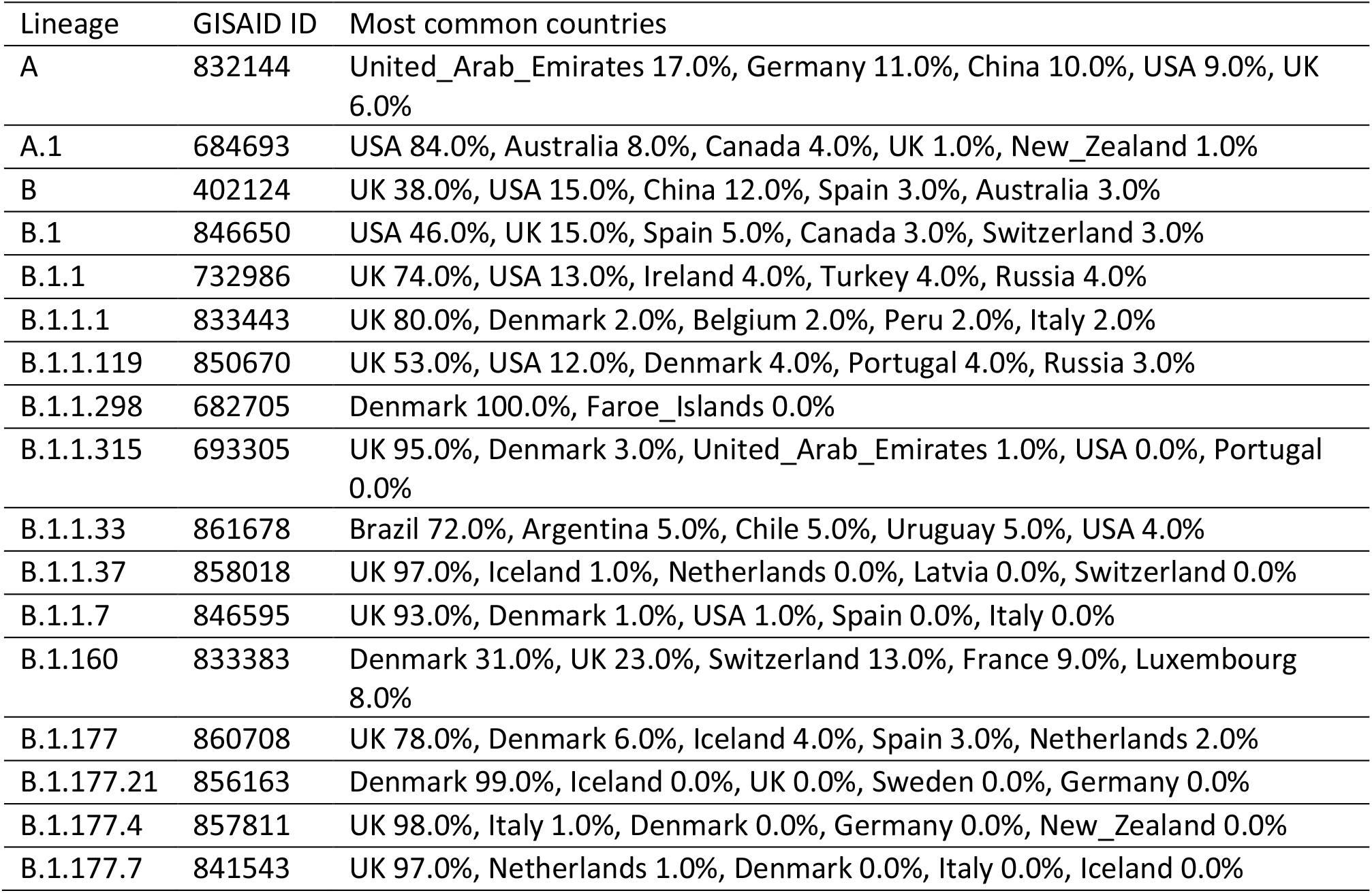
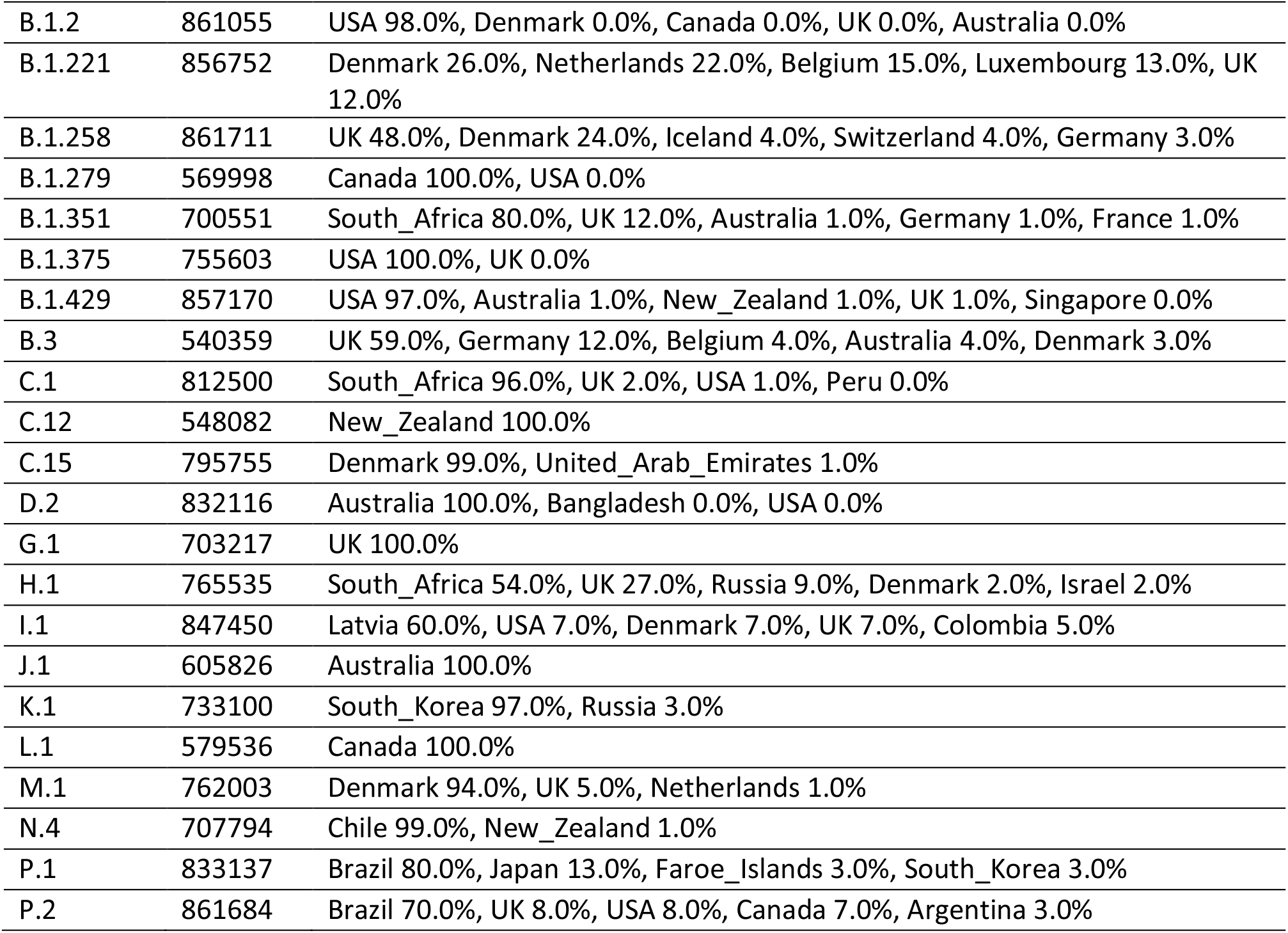
The complete SARS-CoV-2 genomes used for the oligonucleotide design

| Lineage | GISAID ID | Most common countries |
| --- | --- | --- |
| A | 832144 | United_Arab_Emirates 17.0%, Germany 11.0%, China 10.0%, USA 9.0%, UK 6.0% |
| A.1 | 684693 | USA 84.0%, Australia 8.0%, Canada 4.0%, UK 1.0%, New_Zealand 1.0% |
| B | 402124 | UK 38.0%, USA 15.0%, China 12.0%, Spain 3.0%, Australia 3.0% |
| B.1 | 846650 | USA 46.0%, UK 15.0%, Spain 5.0%, Canada 3.0%, Switzerland 3.0% |
| B.1.1 | 732986 | UK 74.0%, USA 13.0%, Ireland 4.0%, Turkey 4.0%, Russia 4.0% |
| B.1.1.1 | 833443 | UK 80.0%, Denmark 2.0%, Belgium 2.0%, Peru 2.0%, Italy 2.0% |
| B.1.1.119 | 850670 | UK 53.0%, USA 12.0%, Denmark 4.0%, Portugal 4.0%, Russia 3.0% |
| B.1.1.298 | 682705 | Denmark 100.0%, Faroe_Islands 0.0% |
| B.1.1.315 | 693305 | UK 95.0%, Denmark 3.0%, United_Arab_Emirates 1.0%, USA 0.0%, Portugal 0.0% |
| B.1.1.33 | 861678 | Brazil 72.0%, Argentina 5.0%, Chile 5.0%, Uruguay 5.0%, USA 4.0% |
| B.1.1.37 | 858018 | UK 97.0%, Iceland 1.0%, Netherlands 0.0%, Latvia 0.0%, Switzerland 0.0% |
| B.1.1.7 | 846595 | UK 93.0%, Denmark 1.0%, USA 1.0%, Spain 0.0%, Italy 0.0% |
| B.1.160 | 833383 | Denmark 31.0%, UK 23.0%, Switzerland 13.0%, France 9.0%, Luxembourg 8.0% |
| B.1.177 | 860708 | UK 78.0%, Denmark 6.0%, Iceland 4.0%, Spain 3.0%, Netherlands 2.0% |
| B.1.177.21 | 856163 | Denmark 99.0%, Iceland 0.0%, UK 0.0%, Sweden 0.0%, Germany 0.0% |
| B.1.177.4 | 857811 | UK 98.0%, Italy 1.0%, Denmark 0.0%, Germany 0.0%, New_Zealand 0.0% |
| B.1.177.7 | 841543 | UK 97.0%, Netherlands 1.0%, Denmark 0.0%, Italy 0.0%, Iceland 0.0% |
| B.1.2 | 861055 | USA 98.0%, Denmark 0.0%, Canada 0.0%, UK 0.0%, Australia 0.0% |
| B.1.221 | 856752 | Denmark 26.0%, Netherlands 22.0%, Belgium 15.0%, Luxembourg 13.0%, UK 12.0% |
| B.1.258 | 861711 | UK 48.0%, Denmark 24.0%, Iceland 4.0%, Switzerland 4.0%, Germany 3.0% |
| B.1.279 | 569998 | Canada 100.0%, USA 0.0% |
| B.1.351 | 700551 | South_Africa 80.0%, UK 12.0%, Australia 1.0%, Germany 1.0%, France 1.0% |
| B.1.375 | 755603 | USA 100.0%, UK 0.0% |
| B.1.429 | 857170 | USA 97.0%, Australia 1.0%, New_Zealand 1.0%, UK 1.0%, Singapore 0.0% |
| B.3 | 540359 | UK 59.0%, Germany 12.0%, Belgium 4.0%, Australia 4.0%, Denmark 3.0% |
| C.1 | 812500 | South_Africa 96.0%, UK 2.0%, USA 1.0%, Peru 0.0% |
| C.12 | 548082 | New_Zealand 100.0% |
| C.15 | 795755 | Denmark 99.0%, United_Arab_Emirates 1.0% |
| D.2 | 832116 | Australia 100.0%, Bangladesh 0.0%, USA 0.0% |
| G.1 | 703217 | UK 100.0% |
| H.1 | 765535 | South_Africa 54.0%, UK 27.0%, Russia 9.0%, Denmark 2.0%, Israel 2.0% |
| I.1 | 847450 | Latvia 60.0%, USA 7.0%, Denmark 7.0%, UK 7.0%, Colombia 5.0% |
| J.1 | 605826 | Australia 100.0% |
| K.1 | 733100 | South_Korea 97.0%, Russia 3.0% |
| L.1 | 579536 | Canada 100.0% |
| M.1 | 762003 | Denmark 94.0%, UK 5.0%, Netherlands 1.0% |
| N.4 | 707794 | Chile 99.0%, New_Zealand 1.0% |
| P.1 | 833137 | Brazil 80.0%, Japan 13.0%, Faroe_Islands 3.0%, South_Korea 3.0% |
| P.2 | 861684 | Brazil 70.0%, UK 8.0%, USA 8.0%, Canada 7.0%, Argentina 3.0% |

Regardless of the technology used for the nucleic acid (NA) extraction, the Bio-Speedy^®^ SARS-CoV-2 N501Y Mutation Detection Kit is applied to NA extracts previously shown to be RT-qPCR positive for SARS-CoV-2 with a threshold cycle (Cq) less than 32. Orf1ab target in the kit ensures that the NA extract kept its integrity during its storage at the temperature recommended by manufacturer of the NA extraction kit. The NA templates with Cq values more than 32 for the Orf1ab target may result in false-negative results since the LOD of the Spike-501 targeted oligo sets are 10x worse than that of the Orf1ab target. The NA extracts must be transferred in dry ice.

The qPCR instruments and their software validated with the Bio-Speedy^®^ SARS-CoV-2 N501Y Mutation Detection Kit is given in Table 4. The details of the reaction set-up and the qPCR program is given in Table 5. The recommended threshold level to calculate the number of threshold cycles (Cq) is 200 RFU for Insta Q96^®^ Plus and CFX96 Touch™ instruments. The threshold level is 0.05 for Roche LightCycler^®^ 96. “Dynamic Tube” should be active, “Slope Correct” should be passive, “Outlier Removal” should be “0”, and the threshold level should be set to 0.02 to calculate the Cq values in Rotor-Gene^®^ Q. For BMS MIC qPCR, “Non-Assay Green/Parameters/Fixed Length” options should be selected, auto-threshold setting should be active.

**Table 4.**
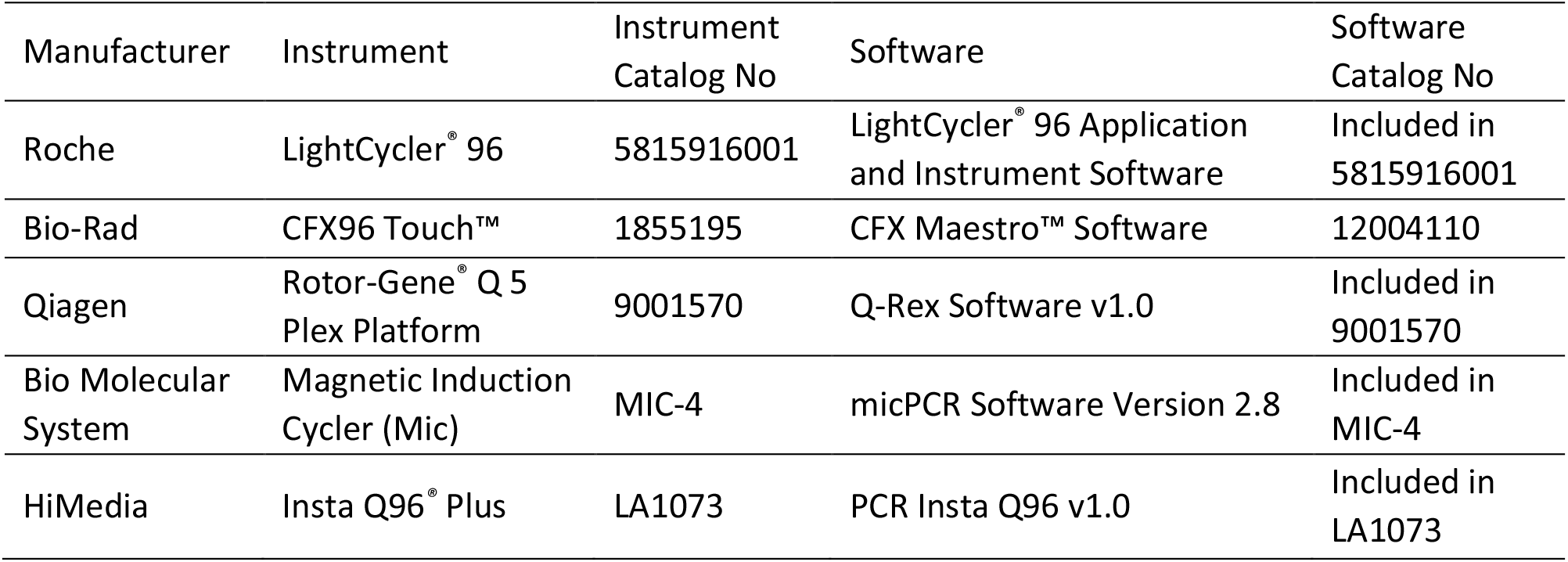
Instruments and their software validated with the Bio-Speedy^®^ SARS-CoV-2 N501Y Mutation Detection Kit

**Table 5.**
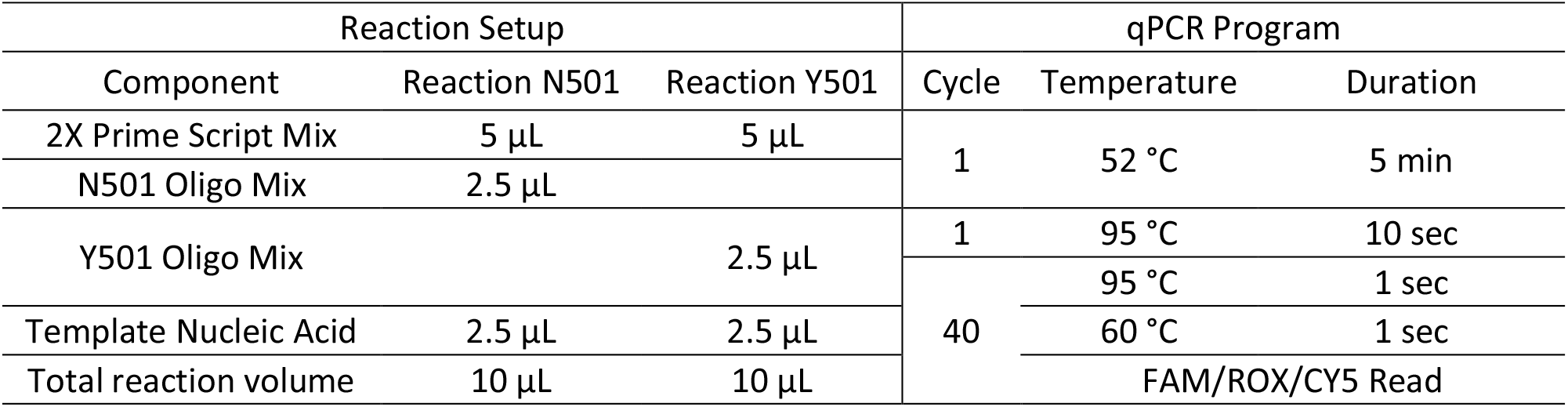
Reaction set-up and the details of the qPCR program

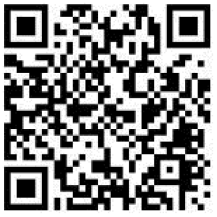

The shape of the amplification curves should be examined. If a Cq value is assigned to a sample by the instruments’ software and the curve is sigmoidal, the Cq value can be used in the final evaluation. Non-sigmoidal curves should be recorded as negative. If a Cq value is assigned to a sample, but the curve is not sigmoidal, the result should be recorded as negative. On the web page linked with the QR code, examples of the sigmoidal amplification curves are given. The results obtained with the Bio-Speedy^®^ SARS-CoV-2 N501Y Mutation Detection Kit should not be interpreted without examining these samples.

If the IC does not perform as described in Table 6, repeat the analysis by increasing the reaction volume to 50 µL by keeping the ratios of the reaction components in Table 5. If the problem continues, then conclude as an invalid PCR template.

**Table 6.** Expected performance of the Bio-Speedy^®^ SARS-CoV-2 N501Y Mutation Detection Kit controls

| Control Type | Name | Control purpose | Expected Result |  |  |
| --- | --- | --- | --- | --- | --- |
|  |  |  | N/Y501 (FAM) | ORF1ab (ROX) | HV69-70del (CY5) |
| Negative Template Control | NTC | Contamination | No Cq (Negative) |  |  |
| Positive Control | PC | Reagent integrity | Cq<38.0 (Positive) |  |  |
| Internal Control | IC | SARS-CoV-2 RNA integrity | Not applicable | Cq<32.0 (Positive) | Not applicable |

If the IC perform as described in Table 6, but there is no amplification in both the N501 and Y501 reactions, repeat the analysis by decreasing the annealing temperature from 60 °C to 55 °C during the 40 cycles of the amplification step (Table 5).

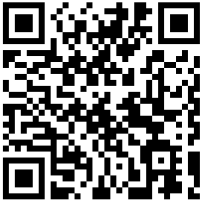

If all the controls are valid, proceed to the interpretation of the results. The “N501 Oligo Mix” targets a SARS-CoV-2 reference genome region encoding the N amino acid at position 501 (**FAM**) of the S protein. The “Y501 Oligo Mix” targets a VOC-202012/01 genome region encoding the Y amino acid at position 501 (**FAM**) of the S protein. There is a single nucleotide difference between the S-501-target of the N501 and Y501 oligo mixes. Hence, the “Y501 Oligo Mix” must amplify Spike-501-target better (with lower Cq values) than the “N501 Oligo Mix” does to obtain the positive N501Y mutation result. The differentiation is made by calculating the Cq value difference between the FAM and ROX channels for each reaction, and by comparing the Cq value differences in the two reactions. Cq difference between Y501 (**FAM**) and Orf1ab (**ROX**) channels in “Y501 Oligo Mix” reaction must be at least 1 Cq lower than that of the “N501 Oligo Mix” reaction to report N501Y mutation positivity. HV6970del mutation is positive if there is a sigmoidal curve in the (**CY5**) channel and the Cq is lower than 38. The calculation method is given below. On the web page linked with the QR code, an excel document that allows the calculation by entering the Cq values is provided.

[Cq N501 in FAM] – [Cq N501 in ROX] = ΔCq N501; [Cq Y501 in FAM] – [Cq Y501 in ROX] = ΔCq Y501 ΔCq N501 - ΔCq Y501= ΔΔCq 501; N501Y positive if ΔΔCq 501 > 1; N501Y negative if ΔΔCq 501 ≤ 1

## Limit of detection (LOD)

LOD studies determine the lowest detectable concentration of the analyte at which greater or equal to 95% of all (true positive) replicates test positive. A pool of nasopharyngeal, oropharyngeal, and nasal swab samples in Bio-Speedy^®^ vNAT™ Transfer Tube (Cat No: BS-NA-513) that were RT-qPCR negative for SARS-CoV-2 were spiked with a cultured SARS-CoV-2 isolate given in Table 9. Because the vNAT™ buffer lyses SARS-CoV-2 and preserves RNA, the lysate in the vNAT™ Transfer Tube was directly used in RT-qPCR reactions.

**Table 7.** The example results of the RT-qPCR assay

| N501 |  |  | Y501 |  |  | N501Y | HV6970del | VOC-202012/01 | 501Y.V2 or P1 possibility |
| --- | --- | --- | --- | --- | --- | --- | --- | --- | --- |
| FAM | ROX | CY5 | FAM | ROX | CY5 |  |  |  |  |
| 31.3 | 23.6 | 29.8 | 26.2 | 23.8 | 29.6 | Pos | Pos | Pos | No |
| Neg | 25.3 | Neg | 31.2 | 28.1 | Neg | Pos | Neg | Neg | Yes |
| 28.3 | 22.5 | 38.3 | 32.1 | 29.1 | 38.9 | Pos | Neg | Neg | Yes |
| 24.5 | 22.1 | 25.3 | 29.6 | 21.9 | 25.1 | Neg | Pos | Neg | No |
| 29.5 | 26.4 | Neg | Neg | 23.6 | Neg | Neg | Neg | Neg | No |
| 30.4 | 27.4 | Neg | 26.6 | 20.8 | Neg | Neg | Neg | Neg | No |
| Neg | Neg | Neg | 33.1 | 31.6 | Neg | Repeat | Neg | Inconclusive | Inconclusive |
| 27.1 | 28.3 | Neg | Neg | Neg | Neg | Repeat | Neg | Inconclusive | Inconclusive |
| 35.6 | Neg | Neg | 22.5 | 26.4 | Neg | Repeat | Neg | Inconclusive | Inconclusive |
| 27.6 | 24.3 | Neg | 34.2 | Neg | Neg | Repeat | Neg | Inconclusive | Inconclusive |
| Neg | 32.4 | Neg | Neg | 32.6 | Neg | Repeat | Neg | Inconclusive | Inconclusive |
| Neg | Neg | Neg | Neg | Neg | Neg | Repeat | Repeat | Inconclusive | Inconclusive |
| 36.2 | Neg | Neg | 38.1 | Neg | Neg | Repeat | Repeat | Inconclusive | Inconclusive |

**Table 8.** The example comments on the results of the RT-qPCR assay

| Case | Oligo Mix | Channel | Target | Result | Comment 1 | Comment 2 |
| --- | --- | --- | --- | --- | --- | --- |
| Case 1 | N501 | FAM | N501 | + | N501Y=Neg | All SARS-CoV-2 variants containing N501Y mutation are negative |
|  |  | ROX | Orf1ab | + |  |  |
|  |  | CY5 | HV69-70del | - |  |  |
|  | Y501 | FAM | Y501 | - | HV69-70del = Neg |  |
|  |  | ROX | Orf1ab | + |  |  |
|  |  | CY5 | HV69-70del | - |  |  |
| Case 2 | N501 | FAM | N501 | + | N501Y = Neg |  |
|  |  | ROX | Orf1ab | + |  |  |
|  |  | CY5 | HV69-70del | + |  |  |
|  | Y501 | FAM | Y501 | - | HV69-70del = Pos |  |
|  |  | ROX | Orf1ab | + |  |  |
|  |  | CY5 | HV69-70del | + |  |  |
| Case 3 | N501 | FAM | N501 | - | N501Y = Pos | VOC-202012/01 is positive |
|  |  | ROX | Orf1ab | + |  |  |
|  |  | CY5 | HV69-70del | + |  |  |
|  | Y501 | FAM | Y501 | + | HV69-70del = Pos |  |
|  |  | ROX | Orf1ab | + |  |  |
|  |  | CY5 | HV69-70del | + |  |  |
| Case 4 | N501 | FAM | N501 | - | N501Y = Pos | 1- VOC-202012/01 is negative<br>2- 501Y.V2 or P1 may be positive. The sequencing is needed to confirm the E484K and the K417(T/N) coexistence. |
|  |  | ROX | Orf1ab | + |  |  |
|  |  | CY5 | HV69-70del | - |  |  |
|  | Y501 | FAM | Y501 | + | HV69-70del = Neg |  |
|  |  | ROX | Orf1ab | + |  |  |
|  |  | CY5 | HV69-70del | - |  |  |

**Table 9.**
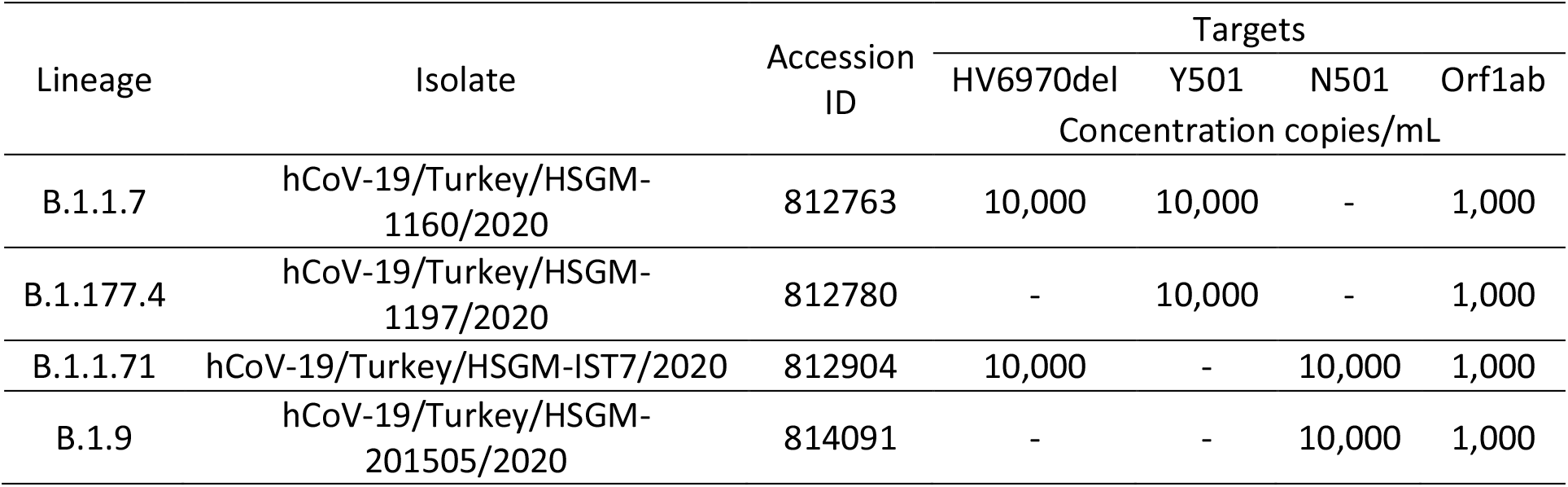
Minimum concentration of the SARS-CoV-2 isolate that was positive in the three replicates of the 10^4^, 10^3^, 10^2^, 10^1^, and 10^0^ copies/mL spiked samples.

The cultured virus was quantified in copies/mL by a molecular assay using 10^3^-10^6^ copies/mL dilutions of synthetic SARS-CoV-2 ORF1ab gene partial synthetic RNA (Oligo-X Biotechnology, Turkey). The acceptance criteria for the dilution series to be used in the standard curves was r^2^>0.9, 1.9<fold-change per cycle<2.0 and percent coefficient of variation (CV%)<5.

Triplicates of the spiked samples with a final concentration of 10^4^, 10^3^, 10^2^, 10^1^, and 10^0^ copies/mL were tested using Bio-Rad CFX96 Touch™ to determine the minimum concentration that yielded positive results in all the replicates. 20 replicates of 2- and 0.5-fold dilutions of the minimum concentration were tested to determine the LOD. 20 replicates of the dilutions at the LOD were tested again using the qPCR instruments in Table 4 other than Bio-Rad CFX96 Touch™.

## Cross-reactivity

A pool of SARS-CoV-2 negative nasopharyngeal, oropharyngeal, and nasal swab samples in the vNAT™ Transfer Tube was spiked with the reference strains and clinical isolates from the culture-confirmed cases. Final concentration of 10^6^ cfu/mL for bacteria and 10^5^ copies/mL for viruses were tested for the cross reactivity. Coronavirus 229E/OC43/NL63/HKU1, MERS, SARS CoV strain Frankfurt 1, Influenza A H1/H3, Influenza B, Parainfluenza 1/2/3/4, Metapneumovirus, Rhinovirus, Respiratory syncytial virus (RSV) A/B, Bocavirus (BoV), Enterovirus, Adenovirus, *Legionella pneumophila, Chlamydia pneumoniae, Mycobacterium tuberculosis, Haemophilus influenzae, Streptococcus pneumoniae, Mycoplasma pneumoniae, Streptococcus pyogenes, Bordetella pertussis, Pneumocystis jirovecii, Candida albicans, Legionella bozemanii, Legionella micdadei, Corynebacterium diphtheriae, Bacillus anthracis, Moraxella catarrhalis, Neisseria meningitidis, Pseudomonas aeruginosa, Staphylococcus epidermidis, Coxiella burneti, Staphylococcus aureus, Streptococcus salivarius, Leptospira interrogans, Chlamydia psittaci* and a pooled nasal wash from 20 different people (healthy donors) were used for testing the exclusivity.

In silico tests were carried out using the BLAST search of the National Center for Biotechnology Information (NCBI). The target region of the oligonucleotide sets was searched against the nucleotide sequence database from which SARS-CoV-2 taxon was excluded.

## Next Generation Sequencing (NGS)

The Bio-Speedy^®^ SARS-CoV-2 N501Y Mutation Detection Kit was applied to 165 clinical samples containing SARS-CoV-2 from 32 different lineages. The lineages were determined via the NGS. 133 SARS-CoV-2 genome sequences with the GISAID accession IDs between 428712-428723, 429861-429873, 811136-811143, 812761-812781, 812873-812921, 814062-814091 were obtained in Illumina MiSeq and assembled using the Burrows-Wheeler Alignerv.07.17-r1188 with 1,000x coverage. 32 SARS-CoV-2 genome sequences with the GISAID accession IDs between 437304 and 437335 were obtained in Oxford Nanopore GridION and assembled using the Geneious Prime 2021.0.3 with 1,000x coverage.

## Sanger Sequencing

The Bio-Speedy^®^ SARS-CoV-2 N501Y Mutation Detection Kit and the Sanger sequencing was applied to 1000 SARS-CoV-2 positive clinical samples collected in Jan2021 from the 81 different provinces of Turkey. The PCR amplicons were obtained using the primer sets shown in Figure 2. The first round was one-step RT-PCR using the 118-F and 1652-R primers that was followed by the nested PCRs using the 118F-695R and 1157F-1652R primers. The Sanger sequencing was performed on Applied Biosystems 3130xl Genetic Analyzer using the reading primers 118F, 641R, 1157F, 1420F and 1652R. The obtained sequences were aligned with the whole SARS-CoV-2 genome sequences obtained via the NGS using the Clustal O tool of the European Bioinformatics Institute (EBI).

**Figure 2.**
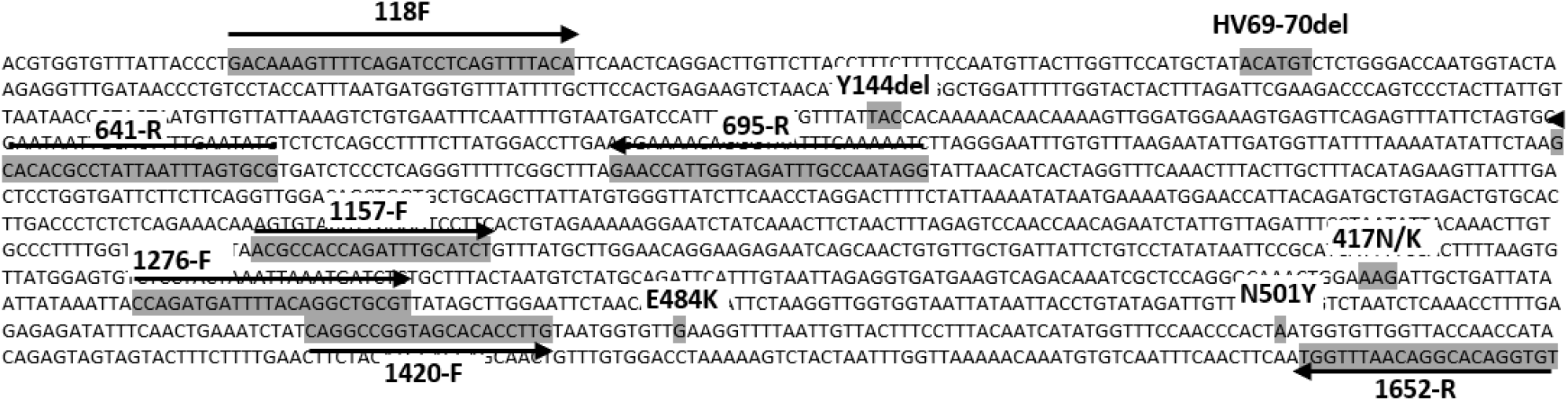
Sanger DNA sequencing primers for Spike N501Y and HV69-70del mutations.

## RESULTS AND DISCUSSION

For the LOD studies given in Table 10-13, the spiked samples were prepared based on the data given in Table 9. The LOD of the Bio-Speedy^®^ SARS-CoV-2 N501Y Mutation Detection Kit for Orf1ab target is 500 copies/mL, while that of the N501, Y501 and HV69-70del targets are 5000 copies/mL.

**Table 10.** LOD of the Bio-Speedy^®^ SARS-CoV-2 N501Y Mutation Detection Kit for the lineage B.1.1.7 in CFX96 Touch™

| Target | Concentration | N501 Oligo Mix |  |  | Y501 Oligo Mix |  |  |
| --- | --- | --- | --- | --- | --- | --- | --- |
|  | copies/mL | Hit Rate | Cq Avg | Cq Std | Hit Rate | Cq Avg | Cq Std |
| Spike N501Y | 20,000 | - | - | - | 20/20 | 34.08 | 0.03 |
|  | 10,000 | - | - | - | 20/20 | 35.12 | 0.04 |
|  | **5,000** | - | - | - | 20/20 | 36.19 | 0.07 |
|  | 2,500 | - | - | - | 14/20 | 37.31 | 0.12 |
|  | 2,000 | - | - | - | 3/20 | 38.44 | 0.56 |
|  | 1,000 | - | - | - | - | - | - |
|  | 500 | - | - | - | - | - | - |
|  | 250 | - | - | - | - | - | - |
| Orf1ab | 20,000 | 20/20 | 31.09 | 0.05 | 20/20 | 31.22 | 0.06 |
|  | 10,000 | 20/20 | 32.11 | 0.04 | 20/20 | 32.26 | 0.08 |
|  | 5,000 | 20/20 | 33.39 | 0.05 | 20/20 | 33.53 | 0.1 |
|  | 2,500 | 20/20 | 34.42 | 0.07 | 20/20 | 34.5 | 0.07 |
|  | 2,000 | 20/20 | 34.83 | 0.07 | 20/20 | 34.87 | 0.08 |
|  | 1,000 | 20/20 | 35.94 | 0.11 | 20/20 | 36.03 | 0.14 |
|  | **500** | 20/20 | 37.32 | 0.19 | 20/20 | 37.46 | 0.24 |
|  | 250 | 16/20 | 38.78 | 0.47 | 14/20 | 38.9 | 0.48 |
| Spike HV-69-70del | 20,000 | 20/20 | 33.82 | 0.02 | 20/20 | 33.96 | 0.04 |
|  | 10,000 | 20/20 | 34.85 | 0.05 | 20/20 | 34.98 | 0.06 |
|  | **5,000** | 20/20 | 36.01 | 0.07 | 20/20 | 36.1 | 0.1 |
|  | 2,500 | 15/20 | 37.23 | 0.12 | 13/20 | 37.27 | 0.13 |
|  | 2,000 | 6/20 | 38.21 | 0.57 | 4/20 | 38.36 | 0.61 |
|  | 1,000 | - | - | - | - | - | - |
|  | 500 | - | - | - | - | - | - |
|  | 250 | - | - | - | - | - | - |

The melting temperature (Tm) of the sense primers targeting the mutations was kept low to increase the effect of the sequence mismatches on the annealing. This resulted in 10x lower LOD and at least 3 cycles lower Cq values of the mutation targets compared to those of the Orf1ab target. Hence, the developed assay should be applied to NA extracts previously shown to be RT-qPCR positive for SARS-CoV-2 with a Cq less than 32 to prevent negative results in the FAM channels of both the N501 and Y501 oligo mixes.

The developed RT-qPCR assay was negative for all strains used in the exclusion tests. The in-silico tests also revealed that the oligonucleotide sets of the assay did not cross-react any nucleotide sequence in the database.

The Bio-Speedy^®^ SARS-CoV-2 N501Y Mutation Detection Kit was tested on 165 SARS-CoV-2 positive samples from which the SARS-CoV-2 whole genomes were sequenced by the Ministry of Health Turkey (Table 14). The RT-qPCR results were in 100% agreement with the whole genome sequence data. 19 of the 29 N501Y positive samples were VOC-202012/01 all detected in people who have traveled from England to Turkey.

**Table 11.** LOD of the Bio-Speedy^®^ SARS-CoV-2 N501Y Mutation Detection Kit for the lineage B.1.1.71 in CFX96 Touch™

| Target | Concentration | N501 Oligo Mix |  |  | Y501 Oligo Mix |  |  |
| --- | --- | --- | --- | --- | --- | --- | --- |
|  | copies/mL | Hit Rate | Cq Avg | Cq Std | Hit Rate | Cq Avg | Cq Std |
| Spike N501Y | 20,000 | 20/20 | 34.12 | 0.05 | - | - | - |
|  | 10,000 | 20/20 | 35.18 | 0.04 | - | - | - |
|  | **5,000** | 20/20 | 36.26 | 0.11 | - | - | - |
|  | 2,500 | 13/20 | 37.39 | 0.09 | - | - | - |
|  | 2,000 | 4/20 | 38.46 | 0.53 | - | - | - |
|  | 1,000 | - | - | - | - | - | - |
|  | 500 | - | - | - | - | - | - |
|  | 250 | - | - | - | - | - | - |
| Orf1ab | 20,000 | 20/20 | 31.18 | 0.01 | 20/20 | 31.08 | 0.1 |
|  | 10,000 | 20/20 | 32.19 | 0.06 | 20/20 | 32.11 | 0.18 |
|  | 5,000 | 20/20 | 33.52 | 0.02 | 20/20 | 33.49 | 0.13 |
|  | 2,500 | 20/20 | 34.54 | 0.06 | 20/20 | 34.38 | 0.08 |
|  | 2,000 | 20/20 | 34.97 | 0.03 | 20/20 | 34.73 | 0.12 |
|  | 1,000 | 20/20 | 35.98 | 0.08 | 20/20 | 35.95 | 0.17 |
|  | **500** | 20/20 | 37.46 | 0.18 | 20/20 | 37.33 | 0.25 |
|  | 250 | 14/20 | 38.93 | 0.47 | 15/20 | 38.81 | 0.48 |
| Spike HV-69-70del | 20,000 | 20/20 | 33.7 | 0.05 | 20/20 | 33.82 | 0.07 |
|  | 10,000 | 20/20 | 34.77 | 0.07 | 20/20 | 34.83 | 0.07 |
|  | **5,000** | 20/20 | 35.88 | 0.07 | 20/20 | 36.06 | 0.11 |
|  | 2,500 | 14/20 | 37.14 | 0.13 | 14/20 | 37.13 | 0.15 |
|  | 2,000 | 6/20 | 38.07 | 0.58 | 4/20 | 38.23 | 0.64 |
|  | 1,000 | - | - | - | - | - | - |
|  | 500 | - | - | - | - | - | - |
|  | 250 | - | - | - | - | - | - |

**Table 12.**
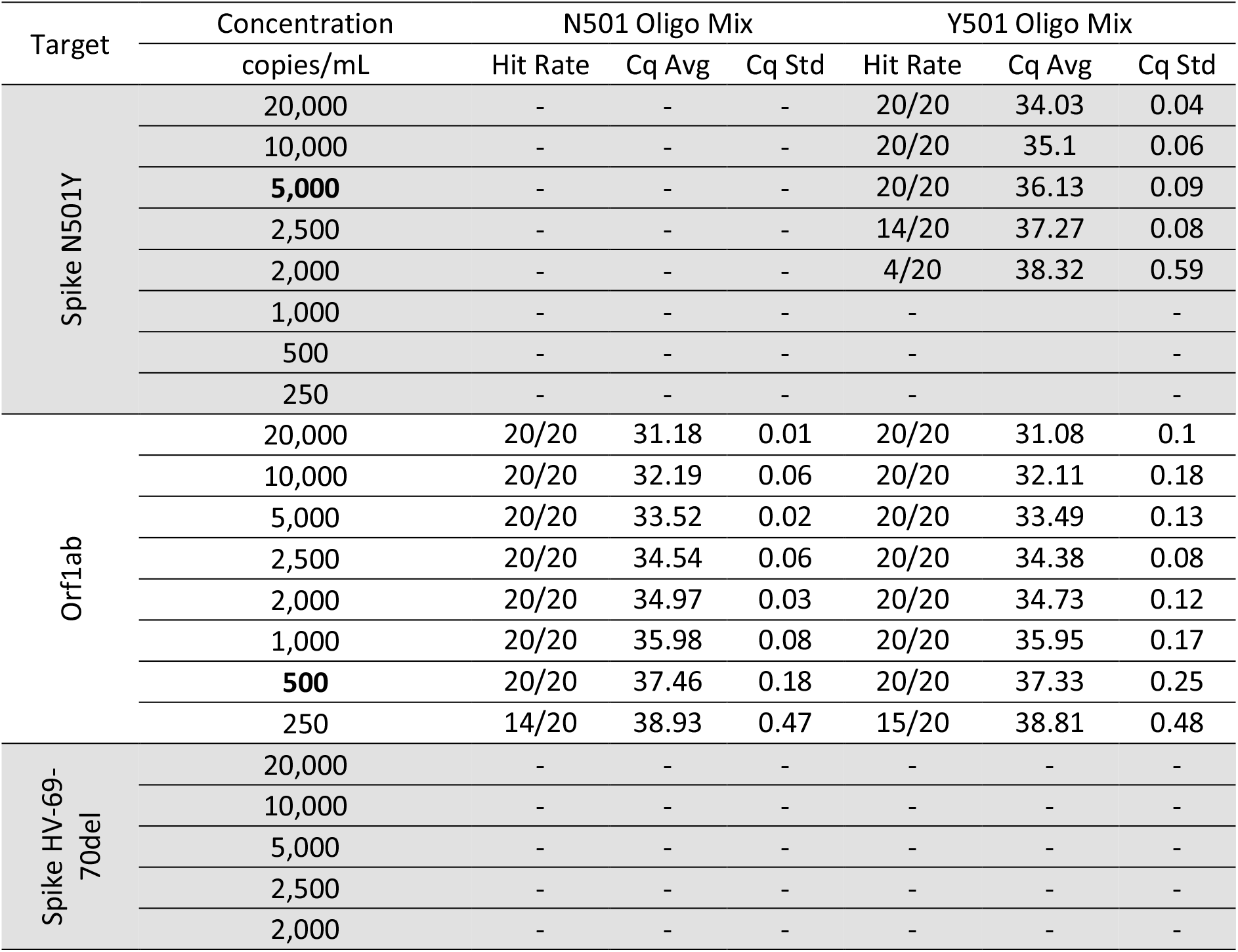
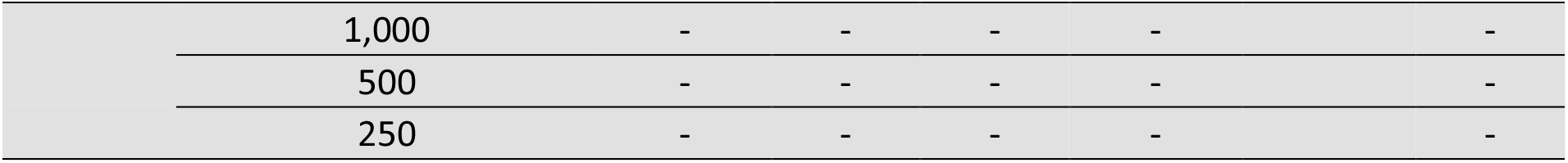
LOD of the Bio-Speedy^®^ SARS-CoV-2 N501Y Mutation Detection Kit for the lineage B.1.177.4 in CFX96 Touch™

| Target | Concentration | N501 Oligo Mix |  |  | Y501 Oligo Mix |  |  |
| --- | --- | --- | --- | --- | --- | --- | --- |
|  | copies/mL | Hit Rate | Cq Avg | Cq Std | Hit Rate | Cq Avg | Cq Std |
| Spike N501Y | 20,000 | - | - | - | 20/20 | 34.03 | 0.04 |
|  | 10,000 | - | - | - | 20/20 | 35.1 | 0.06 |
|  | **5,000** | - | - | - | 20/20 | 36.13 | 0.09 |
|  | 2,500 | - | - | - | 14/20 | 37.27 | 0.08 |
|  | 2,000 | - | - | - | 4/20 | 38.32 | 0.59 |
|  | 1,000 | - | - | - | - | - | - |
|  | 500 | - | - | - | - | - | - |
|  | 250 | - | - | - | - | - | - |
| Orf1ab | 20,000 | 20/20 | 31.18 | 0.01 | 20/20 | 31.08 | 0.1 |
|  | 10,000 | 20/20 | 32.19 | 0.06 | 20/20 | 32.11 | 0.18 |
|  | 5,000 | 20/20 | 33.52 | 0.02 | 20/20 | 33.49 | 0.13 |
|  | 2,500 | 20/20 | 34.54 | 0.06 | 20/20 | 34.38 | 0.08 |
|  | 2,000 | 20/20 | 34.97 | 0.03 | 20/20 | 34.73 | 0.12 |
|  | 1,000 | 20/20 | 35.98 | 0.08 | 20/20 | 35.95 | 0.17 |
|  | **500** | 20/20 | 37.46 | 0.18 | 20/20 | 37.33 | 0.25 |
|  | 250 | 14/20 | 38.93 | 0.47 | 15/20 | 38.81 | 0.48 |
| Spike HV-69-70del | 20,000 | - | - | - | - | - | - |
|  | 10,000 | - | - | - | - | - | - |
|  | 5,000 | - | - | - | - | - | - |
|  | 2,500 | - | - | - | - | - | - |
|  | 2,000 | - | - | - | - | - | - |

|  |  |  |  |  |  |  |
| --- | --- | --- | --- | --- | --- | --- |
|  | 1,000 | - | - | - | - | - |
|  | 500 | - | - | - | - | - |
|  | 250 | - | - | - | - | - |

**Table 13.** LOD of the Bio-Speedy^®^ SARS-CoV-2 N501Y Mutation Detection Kit for the lineage B.1.9 in CFX96 Touch™

| Target | Concentration | N501 Oligo Mix |  |  | Y501 Oligo Mix |  |  |
| --- | --- | --- | --- | --- | --- | --- | --- |
|  | copies/mL | Hit Rate | Cq Avg | Cq Std | Hit Rate | Cq Avg | Cq Std |
| Spike N501Y | 20,000 | 20/20 | 34.09 | 0.03 | - | - | - |
|  | 10,000 | 20/20 | 35.11 | 0.09 | - | - | - |
|  | **5,000** | 20/20 | 36.14 | 0.05 | - | - | - |
|  | 2,500 | 13/20 | 37.31 | 0.09 | - | - | - |
|  | 2,000 | 5/20 | 38.4 | 0.63 | - | - | - |
|  | 1,000 | - | - | - | - | - | - |
|  | 500 | - | - | - | - | - | - |
|  | 250 | - | - | - | - | - | - |
| Orf1ab | 20,000 | 20/20 | 31.23 | 0.02 | 20/20 | 31.11 | 0.11 |
|  | 10,000 | 20/20 | 32.12 | 0.05 | 20/20 | 32.16 | 0.08 |
|  | 5,000 | 20/20 | 33.48 | 0.01 | 20/20 | 33.51 | 0.1 |
|  | 2,500 | 20/20 | 34.52 | 0.06 | 20/20 | 34.32 | 0.08 |
|  | 2,000 | 20/20 | 34.91 | 0.04 | 20/20 | 34.78 | 0.13 |
|  | 1,000 | 20/20 | 36.03 | 0.08 | 20/20 | 36.09 | 0.19 |
|  | **500** | 20/20 | 37.54 | 0.16 | 20/20 | 37.28 | 0.24 |
|  | 250 | 13/20 | 38.89 | 0.52 | 12/20 | 38.74 | 0.51 |
| Spike HV-69-70del | 20,000 | - | - | - | - | - | - |
|  | 10,000 | - | - | - | - | - | - |
|  | 5,000 | - | - | - | - | - | - |
|  | 2,500 | - | - | - | - | - | - |
|  | 2,000 | - | - | - | - | - | - |
|  | 1,000 | - | - | - | - | - | - |
|  | 500 | - | - | - | - | - | - |
|  | 250 | - | - | - | - | - | - |

**Table 14.** Lineages of the SARS-CoV-2 genomes sequenced by the Ministry of Health Turkey (GISAID accession IDs between 428712-428723, 429861-429873, 437304-437335, 811136-811143, 812761-812781, 812873-812921, 814062-814091)

| Lineage | Genotype |  | 2021 |  | 2020 |  |  |  |  |
| --- | --- | --- | --- | --- | --- | --- | --- | --- | --- |
|  | N501Y | HV6970del | Jan | Dec | Sep | Jun | May | Apr | Mar |
| B.1.1.7 | + | + | 5 | 14 | 0 | 0 | 0 | 0 | 0 |
| B.1.1.7 | + | - | 0 | 1 | 0 | 0 | 0 | 0 | 0 |
| B.1.177.4 | + | - | 0 | 2 | 0 | 0 | 0 | 0 | 0 |
| B.1.36 | + | - | 0 | 2 | 0 | 0 | 0 | 0 | 0 |
| A | + | - | 0 | 1 | 0 | 0 | 0 | 0 | 0 |
| B.1.1 | + | - | 1 | 3 | 0 | 0 | 0 | 0 | 0 |
| B.1.1.105 | - | + | 0 | 1 | 0 | 0 | 0 | 0 | 0 |
| B.1.1.105 | - | - | 0 | 1 | 1 | 2 | 1 | 0 | 0 |
| A | - | - | 0 | 2 | 0 | 3 | 0 | 0 | 1 |
| B | - | - | 0 | 0 | 0 | 0 | 0 | 0 | 2 |
| B.1 | - | - | 0 | 10 | 0 | 0 | 0 | 0 | 4 |
| B.1.1 | - | - | 0 | 10 | 1 | 4 | 3 | 1 | 6 |
| B.1.1.10 | - | - | 0 | 0 | 0 | 0 | 0 | 0 | 1 |
| B.1.1.171 | - | - | 0 | 0 | 0 | 0 | 0 | 0 | 1 |
| B.1.1.183 | - | - | 0 | 1 | 0 | 0 | 0 | 0 | 0 |
| B.1.1.189 | - | - | 0 | 6 | 0 | 0 | 0 | 0 | 0 |
| B.1.1.220 | - | - | 0 | 1 | 0 | 0 | 0 | 0 | 0 |
| B.1.1.283 | - | - | 0 | 1 | 0 | 0 | 0 | 0 | 0 |
| B.1.1.288 | - | - | 1 | 4 | 0 | 0 | 1 | 0 | 0 |
| B.1.1.54 | - | - | 0 | 2 | 0 | 0 | 0 | 1 | 0 |
| B.1.126 | - | - | 0 | 0 | 0 | 0 | 0 | 0 | 1 |
| B.1.160 | - | - | 0 | 4 | 0 | 0 | 0 | 0 | 0 |
| B.1.220 | - | - | 0 | 0 | 0 | 0 | 0 | 1 | 0 |
| B.1.246 | - | - | 0 | 1 | 0 | 0 | 0 | 0 | 6 |
| B.1.247 | - | - | 0 | 0 | 0 | 0 | 0 | 0 | 1 |
| B.1.255 | - | - | 0 | 1 | 0 | 0 | 0 | 0 | 0 |
| B.1.272 | - | - | 0 | 0 | 0 | 0 | 0 | 0 | 1 |
| B.1.36 | - | - | 0 | 4 | 0 | 0 | 0 | 0 | 3 |
| B.1.36.13 | - | - | 0 | 0 | 0 | 0 | 0 | 0 | 1 |
| B.1.36.6 | - | - | 0 | 0 | 0 | 2 | 0 | 2 | 3 |
| B.1.5 | - | - | 0 | 1 | 0 | 0 | 1 | 0 | 4 |
| B.1.9 | - | - | 0 | 2 | 1 | 0 | 0 | 1 | 3 |
| B.4 | - | - | 0 | 0 | 0 | 0 | 0 | 0 | 19 |
| 165 SARS-CoV-2 = |  |  | 7 | 75 | 3 | 11 | 6 | 6 | 57 |

The Bio-Speedy^®^ SARS-CoV-2 N501Y Mutation Detection Kit and the Sanger sequencing was applied to 1000 SARS-CoV-2 positive clinical samples collected in Jan2021 from the 81 different provinces of Turkey. The RT-qPCR results were in 100% agreement with the Sanger sequencing results (Table 15).

**Table 15.** Statistics of the sequences obtained using the Sanger method.

| Genotype | Sanger Sequencing<br>Spike aa position |  | Quantity | SARS-CoV-2 strain with 100% sequence homology |  |  |
| --- | --- | --- | --- | --- | --- | --- |
|  | 60-200 | 400-530 |  | Virus name | GISAID ID | Lineage |
| 1 |  | N501Y | 21 | hCoV-19/Turkey/HSGM-1197/2020 | 812780 | B.1.177.4 |
| 2 |  | N501Y | 8 | hCoV-19/Turkey/HSGM-1125/2020 | 812776 | B.1.36 |
| 3 |  | N501Y | 3 | hCoV-19/Turkey/HSGM-1192/2020 | 811143 | B.1.1.189 |
| 4 | HV69-70del, V171F |  | 4 | hCoV-19/Turkey/HSGM-IST7/2020 | 812904 | B.1.1.105 |
| 5 | G181R |  | 128 | hCoV-19/Turkey/HSGM-IST23/2020 | 812919 | B.1.36 |
| 6 |  | D467Y | 92 | hCoV-19/Turkey/HSGM-2002084/2020 | 814090 | B.1.1.189 |
| 7 |  | S477N | 87 | hCoV-19/Turkey/HSGM-AD1/2020 | 812886 | B.1.160 |
| 8 |  | A520S | 72 | hCoV-19/Turkey/HSGM-IST5/2020 | 812902 | B.1.1.102 |
| 9 |  | I468V | 68 | hCoV-19/Turkey/HSGM-1009/2020 | 437322 | B.4 |
| 10 | D138H |  | 97 | hCoV-19/Turkey/HSGM-AD13/2020 | 812894 | B.1.1.54 |
| 11 | A93S, K97N |  | 74 | hCoV-19/Turkey/HSGM-IST20/2020 | 812916 | B.1.1.54 |
| 12 |  | D467Y, S469A | 63 | hCoV-19/Turkey/HSGM-2002084/2020 | 814090 | B.1.1.189 |
| 13 | V70F | A520S | 139 | hCoV-19/Turkey/HSGM-IST10/2020 | 812906 | B.1.1.189 |
| 14 | SG71-72del | N501T | 18 | hCoV-19/Turkey/HSGM-AD17/2020 | 812897 | A |
| 15 |  | Y508H | 126 | hCoV-19/Turkey/HSGM-19634/2020 | 812873 | B.1.1.215 |

Increasing trend of the VOC-202012/01 prevalence is the indicator of its higher transmission rates. The 501Y.V2 and P.1 variants do not seem to have a higher transmission rate, but they are also alarming due to their potential of causing more severe illness or escaping from the neutralizing antibodies. The developed RT-qPCR assay allows detection of both VOC-202012/01 and the suspected cases of the 501Y.V2 and P.1 variants with considerably lower time and cost compared to the sequencing-based technologies.

The NGS is still the only way to detect emergence of a new and threatening SARS-CoV-2 variant, but the RT-qPCR is the best tool to track prevalence of the new threat in a timely and representative manner.

Currently, VOC-202012/01 is spreading globally, like the SARS-CoV-2 variant with the D614G mutation became dominant in the first quarter of 2020. (13). The UK experience taught us that if the VOC-202012/01 abundance is increasing even though its prevalence is low, a massive increase in the SARS-CoV-2 positive cases would be expected in the future. Hence, entry of the VOC-202012/01 carriers should be prevented at the country borders, filiation studies should be accelerated for the identified cases and more specific quarantine conditions should be created. This is possible only by testing the highest portion of the positives in the shortest time. The developed 40 minutes RT-qPCR assay allows this because the quantity of the qPCR instruments has been increasing tremendously while its application cost is decreasing all around the world.

